# Quantifying evolutionary novelty and design efficiency in generative genome design

**DOI:** 10.64898/2026.06.12.731871

**Authors:** James R.M. Black, Aaron Maiwald, Jassi Pannu, Oliver M. Crook

## Abstract

Generative genome design models can now produce previously unobserved genome-length sequences, but assessing their capabilities is complicated by limitations in functional prediction. The ability to engineer genomes faster than we can understand them risks creating biosecurity vulnerabilities. To evaluate these potential risks systematically, we propose a framework that distinguishes between (i) *evolutionary novelty*, quantified through phylogenetic and sequence similarity to natural genomes; and (ii) *design efficiency* - the efficiency with which a model finds viable sequences compared to simple baseline generators. Applying this framework to bacteriophages designed by the genome language model Evo 2, we find that model likelihood strongly predicts experimental viability, capturing functional constraints beyond simple biological heuristics. However, this efficiency derives largely from staying close to previously observed sequences rather than exploring novel sequence space, reflecting the combined performance of the model and additional filters that were applied to its outputs. Compared to baselines of random mutagenesis and serial passage, the model achieves substantial design efficiency while its outputs remain phylogenetically close to natural genomes. We conclude that the generative capabilities of Evo 2 warrant low to moderate biosecurity concern for de novo hazard creation, although the degree to which these findings generalise to larger or less constrained viral architectures is an open question. Our framework enables an evidence-based capability assessment of generative genome design tools, informing future biosecurity evaluations.

## 1 Introduction

Genome language models have evolved from early predictive models (Ji et al., 2021) to powerful generative tools capable of designing genes and whole genomes that have been validated *in vitro* (Nguyen et al., 2024; Brixi et al., 2025). These generative capabilities follow rapid progress in models’ predictive performance. This includes predicting the effects of genetic variants in plant and vertebrate genomes (Benegas et al., 2023, 2025; Mendoza-Revilla et al., 2024; Zhai et al., 2025), annotating microbial or plant genes (Jha et al., 2024; Almeida et al., 2025), and identifying regulatory elements (Schiff et al., 2024; Fishman et al., 2025; Dalla-Torre et al., 2025). These predictive capabilities have the potential to accelerate scientific discovery in genomics, improve diagnostic yield for genetic disease and enable low-N prioritisation of variants to reduce the cost of wet-lab validation.

Characterising the generative capabilities of these models is a critical challenge. Genome language models can generate diverse sequences by adjusting sampling parameters (such as temperature), but increased diversity typically comes at the cost of biological viability. Since *in silico* functional prediction remains unreliable for novel sequences, our understanding of model capabilities is likely to continue to lag behind their raw generative capacity. Empirical *in vitro* validation therefore remains the gold standard for benchmarking, and has been specifically recommended for biosecurity assessment (Bloomfield et al., 2024; Pannu et al., 2025).

A recent paper evaluated the viability of synthetic bacteriophage designs produced by Evo 2, a state-of-the-art genome language model (King et al., 2025). The bacteriophage chosen by the authors as a model, Phi X-174, is well-suited for synthetic genome design validation: it has a small, modular genome, its short replication cycle enables high-throughput testing, and its inability to infect humans offers significant safety advantages over human-tropic viruses (Hatfull & Hendrix, 2011; Kilcher & Loessner, 2019; Ranveer et al., 2024). This dataset provides the first opportunity to assess generative genome design capabilities against real functional validation data.

Evaluating generative capabilities is complicated by the conflation of two distinct concepts: sequence novelty and design efficiency. These require separate treatment: a model might efficiently identify viable sequences while remaining close to known biology, or explore radically novel space while rarely producing functional outputs. To disentangle these, we propose a two-axis framework that separates (a) the novelty of sequences a model can generate from (b) the efficiency with which it identifies viable candidates.

The first axis captures evolutionary novelty: whether model outputs remain within the distribution of natural sequences, themselves a sparse sample of what evolution has explored, or depart from it (see Fig 1A). This distinguishes interpolation within known sequence space from extrapolation beyond it, regardless of whether any particular output is functional. The second axis captures design efficiency: how effectively a model identifies sequences that validate in vitro, compared with baselines like random sampling, rule-based filtering, or directed evolution. A model may be tightly constrained in its exploration yet still provide substantial efficiency gains.

**Figure 1:**
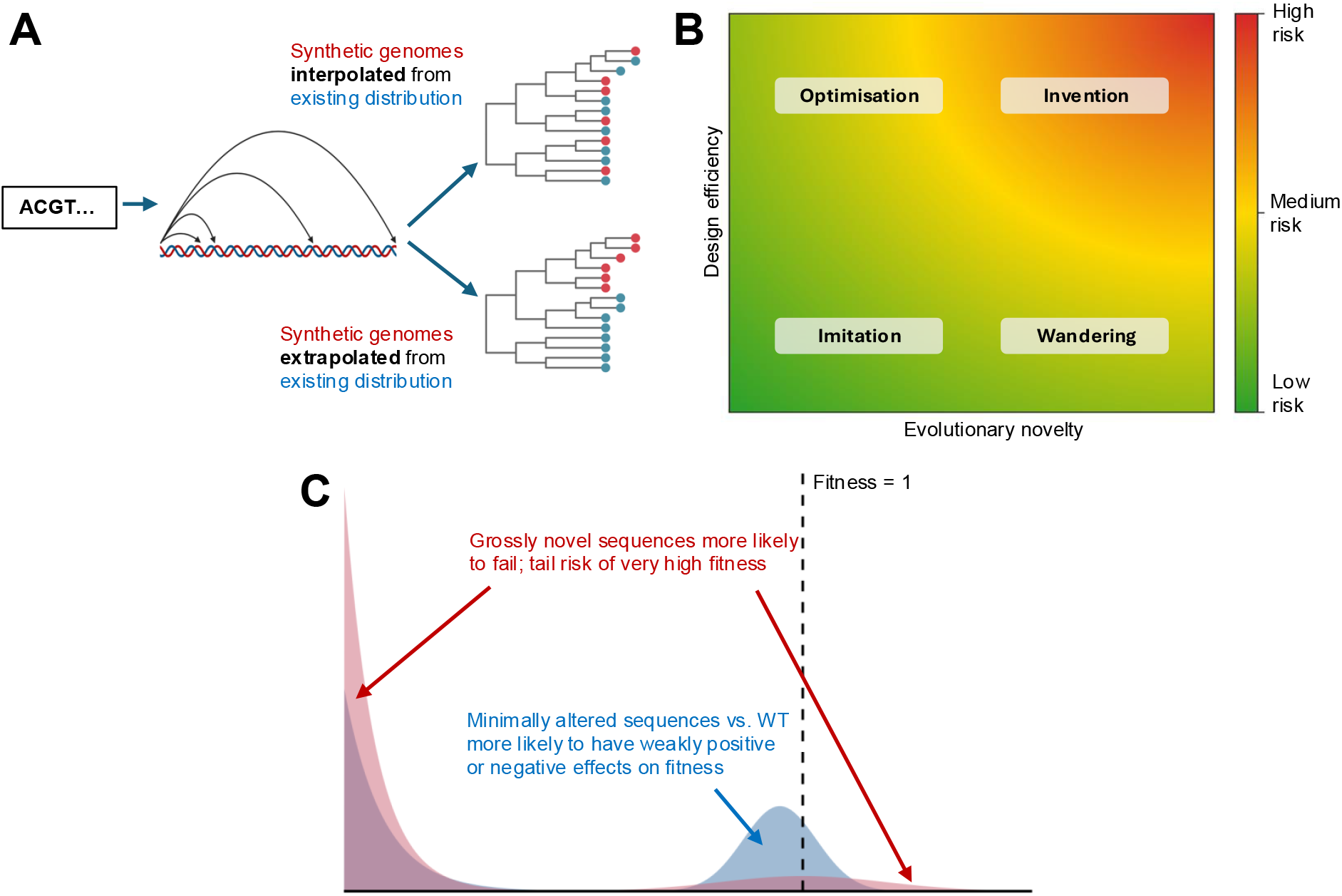
**A)** Generative genome language models use an input nucleotide “prompt” to generate synthetic genomes of varying length. These synthetic genomes can land within the existing distribution of sequences within the training dataset (interpolative) or could theoretically generate fundamentally new clades (extrapolative). **B)** Our framework proposes four categories along the two dimensions of evolutionary novelty and design efficiency, defining four biosecurity risk groups. Putative risk for de novo hazard creation is indicated by colours, with red being the most concerning. **C)** Illustration of hypothesised relationship between the novelty of a modified virus relative to the wild-type and its fitness. Here, modest modifications, such as those conferred in mutagenesis experiments (indicated by the blue shaded area) are more likely to give rise to modest fitness changes than sequences that are highly novel. For sequences of significant novelty (indicated by the red shaded area), such as those that might be created by a generative genome design model, it is more likely that they would be non-viable; however, the probability of extreme outcomes may increase simply because the distribution of functional effects becomes broader.

**Figure 2:**
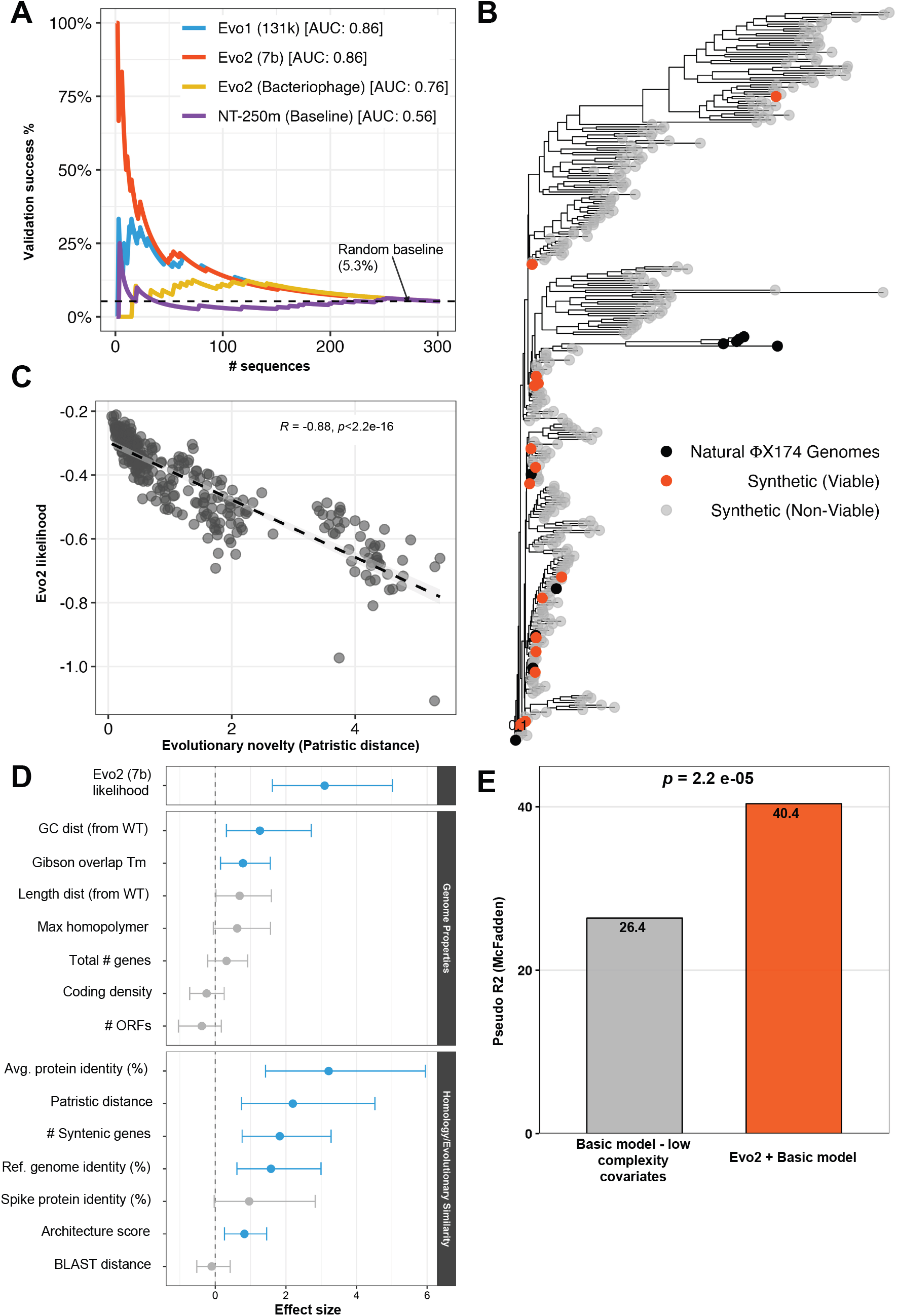
**A)** Sequences generated by King et al. (2025) are ranked according to model probabilities generated from different GLMs and the success rate is plotted as a function of descending through this ranked list. Predictive AUCs are annotated. **B)** A phylogenetic tree of natural ΦX174 genomes along with synthetic viable and non-viable genomes shown. **C)** Evo 2 likelihood correlates strongly with evolutionary novelty for generated sequences. **D)** Forest plot of uni-variable parameters predicting viability from model likelihoods, genome properties and evolution measures. **E)** Pseudo-*R*^2^ for multivariable models using low complexity covariates and a model with low complexity variables in combination with Evo 2 likelihoods.

This distinction matters for biosecurity. The spectrum of concerns about genome language models includes at one end capability enhancement via mutagenesis (predicting more pathogenic variants of existing agents) and, at the other, novel hazard creation (designing altogether novel threats). These map differently onto our framework: capability enhancement relates primarily to design efficiency within known sequence space, while novel hazard creation would require both sufficiently high efficiency to be affordable and exploration beyond evolutionary precedent. Empirical work has begun evaluating predictive capabilities of concern (Black et al., 2025; Wei et al., 2025), but equivalent assessment of generative capabilities is lacking.

Here, we apply this framework to bacteriophage genome design by Evo 2. We find that Evo 2 operates as an evolutionary optimiser: substantially improving design efficiency while producing outputs that remain phylogenetically close to natural sequences. For compact viral genomes, this positions current models as efficiency tools rather than sources of qualitatively novel biological hazards - a distinction with direct implications for biosecurity governance. Of note, the degree to which these conclusions generalise to larger or structurally different viruses is not tested.

## 2 Results

### 2.1 A two-axis framework

To provide empirical evidence to support the degree of biosecurity concern that should be attached to Evo 2 and similar generative genome design models, we designed a two-axis capability assessment framework (Fig 1B). Importantly, this framework applies not to raw model outputs alone, but to the complete model-scaffold: the generative model together with any filtering, ranking, or curation steps applied before sequences are synthesised. In practice, users will apply biological heuristics or use model likelihoods to filter candidates, and it is this combined system that determines real-world capability. Throughout, we refer to the model and the complete model-scaffold interchangeably.

Here, each axis corresponds to a type of behaviour: i) evolutionary novelty: do model-scaffold outputs remain within the statistical distribution of natural sequences, or depart from it?; and ii) design efficiency: to what degree does the model-scaffold identify functional sequences more efficiently than baselines (random sampling, simple heuristics, or directed evolution)? Throughout this work, we analyse evolutionary novelty by sequence divergence from known natural genomes, quantified using phylogenetic distance and sequence similarity metrics. We note that evolutionary novelty in this sense is distinct from functional/biological novelty, since phenotypic innovation may not scale linearly with sequence divergence. For example, a genome that differs from a natural virus by only a small number of mutations could nevertheless acquire a novel host range or other emergent phenotype, whereas highly divergent sequences may remain functionally similar.

This framework naturally suggests four broad categories:

#### 1. Imitation - (Low Novelty, Low Design Efficiency)

Model-scaffolds that produce sequences similar to natural ones, but no more likely to work than simple approaches like random mutagenesis. These systems don’t meaningfully help anyone as they offer no advantage over existing methods.

#### 2. Optimisation - (Low Novelty, High Design Efficiency)

Model-scaffolds that stay close to natural sequences but are substantially better at finding functional ones. For biosecurity, these are powerful tools for accelerating existing workflows: reducing the expertise required or the number of candidates that need to be screened. They make known biology easier to engineer, but don’t enable fundamentally new designs.

#### 3. Wandering - (High Novelty, Low Design Efficiency)

Model-scaffolds that venture into novel sequence space but mostly produce non-functional outputs. The occasional success requires screening many failures. However, such systems could still benefit well-resourced actors with dedicated laboratory capacity willing to screen large numbers of non-functional candidates for rare successes, perhaps to evade screening or detection methods. However, they offer little to actors with limited resources or expertise, for whom high failure rates are prohibitive.

#### 4. Invention - (High Novelty, High Design Efficiency)

Model-scaffolds that reliably produce functional sequences unlike anything seen in nature. Such systems would represent a qualitatively new capability. For biosecurity, this is the primary concern: even actors with modest resources could potentially access novel biological design space. Additionally, actors seeking to evade sequence-based screening, by generating functional sequences with low similarity to known threats, would benefit from this capability regardless of their resource level.

Importantly, individual models can move within this framework depending on sampling parameters (such as temperature) and the stringency of downstream filtering. Higher temperature increases novelty at the cost of viability; stricter filtering improves efficiency but may constrain novelty. Whether high novelty and high efficiency can be jointly achieved, or whether there exists a fundamental tradeoff preventing this, remains an open empirical question. We operationalise novelty through phylogenetic distance and sequence similarity to known genomes, and efficiency through viability rates relative to baseline generators (see Methods). Capability assessments should characterise model-scaffolds across a range of operating conditions rather than at a single point.

### 2.2 Biosecurity concerns of predictive and generative properties of genome language models

Genome language models can operate in two modes. In predictive mode, the model scores existing sequences — useful for ranking mutations or filtering candidates. In generative mode, the model produces novel sequences from scratch. Previous biosecurity evaluations of genome language models have focused on predictive capabilities - the ability to score or rank mutations (Fig 1C, blue distribution). Generative capabilities pose a distinct concern: tolerance for high failure rates in exchange for tail-risk access to highly fit novel sequences (Fig 1C, red distribution). Evaluating this possibility requires functional validation data, which the King et al. (2025) dataset uniquely provides. We therefore assessed both evolutionary novelty and design efficiency of Evo 2’s generative outputs.

### 2.3 Evo 2 likelihoods predict viability better than low-complexity biological covariates

We initially structured our analysis around three questions that link directly to these axes:

1. Does Evo 2 provide design efficiency by identifying viable sequences better than random chance?
2. Does this efficiency extend beyond biological heuristics that one might apply?
3. Does the model achieve this efficiency by staying close to evolutionary precedent or by exploring novel sequence space?

The first two questions address the y-axis (design efficiency); the third addresses the x-axis (evolutionary novelty) and probes the *source* of any observed efficiency.

#### Design efficiency: better than random?

We classified sequences from King et al. (2025) as viable or non-viable based on their *in vitro* validation results (see Methods), then asked whether model likelihoods could discriminate between them. We scored all 302 sequences using four genome language models: Evo 1 (131k context), Evo 2 (7B parameters), the bacteriophage fine-tuned Evo 2 from King et al. (2025), and Nucleotide Transformer (250M parameters) as a non-Evo baseline.

Evo 2 likelihood strongly predicted viability (AUC = 0.86), as did Evo 1 (AUC = 0.86). The fine-tuned model performed worse (AUC = 0.76), indicating that fine-tuning did not improve discrimination. Nucleotide Transformer showed minimal predictive ability (AUC = 0.56), suggesting that design efficiency may be specific to the Evo architecture rather than genome language models generally (Fig. 3A). This confirms substantial design efficiency relative to a random baseline.

**Figure 3:**
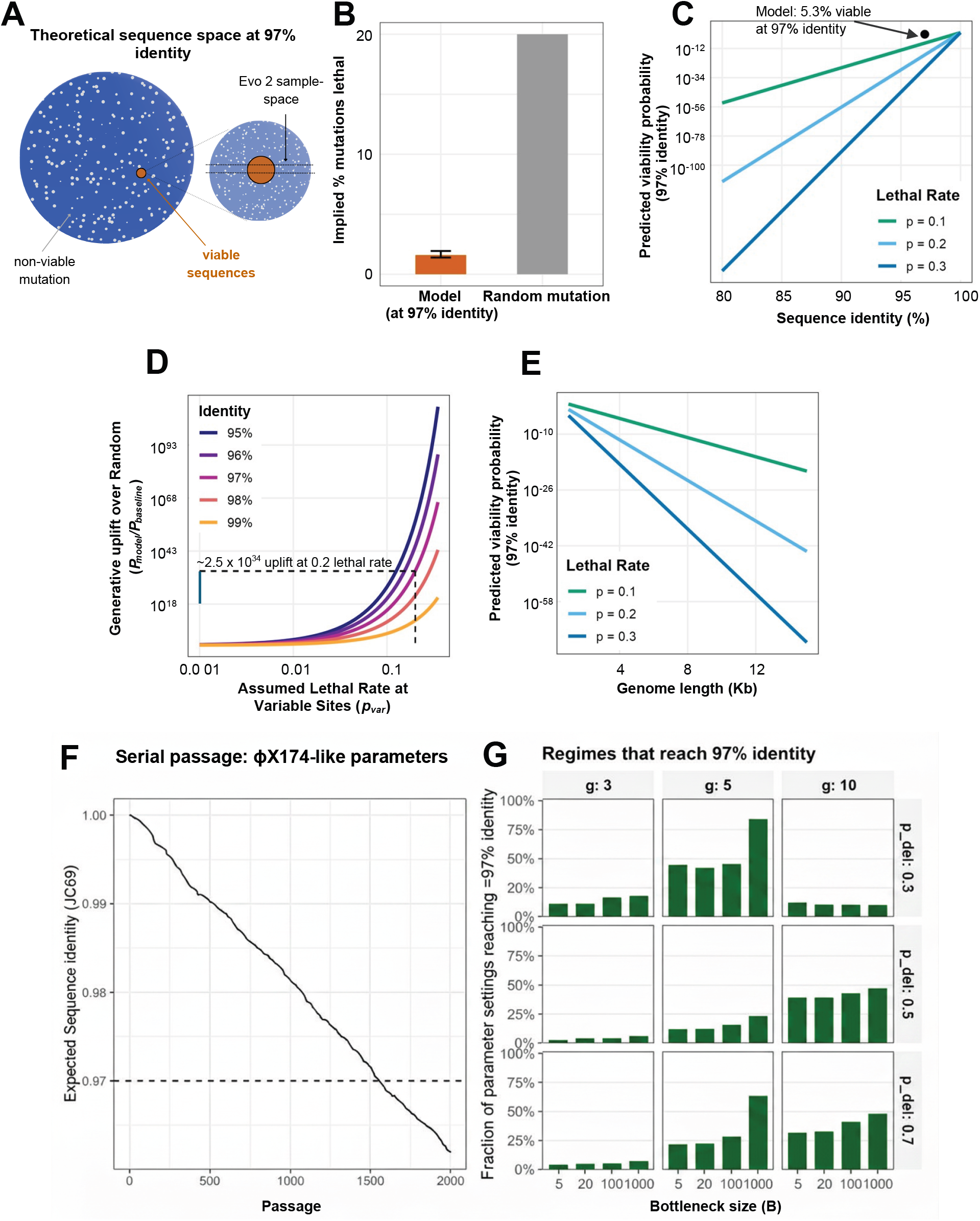
Quantifying design efficiency beyond evolutionary conservation. **A)** Illustrative schematic (not to scale) of sequence space showing the generative model (orange) preferentially targeting the viable manifold relative to random mutagenesis (grey). **B)** Effective per-mutation failure rate (*p*_eff_ *≈* 1.7%) inferred from model viability compared to a baseline of random mutation (*∼* 20%) (Domingo-Calap et al., 2009). **C)** Theoretical probability of sequence being viable is a function of sequence identity; the observed performance of this model (black dot) implies significant uplift relative to a range of plausible rates of lethality. **D)** Generative uplift (*P*_model_*/P*_baseline_) plotted against assumed lethal rates at variable sites, showing 10^30^-fold improvement relative to a MSA-aware baseline. Specific values shown reflect an assumption about the independence of mutations within a genome that is certainly violated within ΦX174. Dashed line represents the per-mutation lethality rate (20%) reported by Domingo-Calap et al. **E)** Projected viability probability as a function of genome length (extrapolated to 15 kb), under the worst-case assumption of uniform mutational constraint throughout the genome. Although putative lethal mutations are likely enriched in small genomes, their absolute number is modelled to scale with genome length, causing the probability of viability via random mutation to decay exponentially (modelled as *P ∝* (1 *− p*)^*L*^). Success probability therefore declines as genomic length increases. F) Example serial passage trajectory with expected sequence identity to the wild-type as a function of passage number under typical parameters. G) Sensitivity analysis of serial passage simulations varying across generation number, bottleneck size and deleterious probability (see Methods)

#### Evolutionary novelty: where does efficiency come from?

High design efficiency could arise in two ways: by learning to navigate novel sequence space effectively (Invention quadrant), or by learning to stay close to sequences that evolution has already validated (Optimisation quadrant). We tested which applies to Evo 2.

We reconstructed a phylogeny from the 302 synthetic sequences and 11 natural ΦX174 reference genomes used by King et al. (2025) (Fig. 3B), selected in order that analyses were directly comparable with the original phylogenetic analyses, and calculated the patristic distance from each synthetic sequence to its nearest natural neighbour (see methods).

Evo 2 likelihood correlated strongly with evolutionary proximity (Pearson’s *ρ* = − 0.88, *p <* 2.2*e*^*−*16^; Fig. 3C): sequences the model scored highly were phylogenetically closer to natural genomes. This indicates that Evo 2’s efficiency derives from recognising and reproducing patterns present in natural variation, rather than from successful extrapolation into unexplored regions of sequence space.

#### Design efficiency: better than heuristics?

A model that merely recapitulates rules a human could specify, such as avoiding GC extremes, preserving gene structure, and maintaining synteny would provide limited added value. We tested whether Evo 2 captures something beyond such heuristics.

We evaluated a suite of low-complexity covariates, including some of those aggregated by King et al.: GC content deviation, assembly thermodynamics, protein similarity, gene synteny, and genomic architecture (Figure 3D; full list in Methods). Several predicted viability in univariate analysis, and together explained 26% of variance (pseudo-R^2^). Adding Evo 2 likelihood significantly improved the model (likelihood ratio *p* = 2.2e-05), increasing variance explained to 40%. Evo 2 therefore captures functional constraints beyond what simple biological rules provide, confirming design efficiency that would be difficult to replicate through curation of these covariates alone (in practice, there may be other structural or regulatory considerations that might not have been captured by this suite of covariates).

### 2.4 Quantifying design efficiency against baseline generators

The previous section established that Evo 2 provides design efficiency, but how much? To place this in a quantitative context, we compared the model’s observed viability rate against baseline generators that represent alternative paths to the same sequence diversity.

#### The challenge of baseline comparison

At 97% nucleotide identity to the reference phage, the model achieved 5.3% success in identifying viabile sequences (16 of 302 sequences). This sounds modest, but the relevant question is: what viability might we expect from alternative approaches operating at the same identity threshold? We considered three baselines:

1. Random mutagenesis: mutations placed at random across the genome
2. MSA-aware mutagenesis: mutations restricted to sites variable in natural alignments
3. Directed evolution: iterative selection over multiple generations

#### Inferring an effective failure rate

To enable comparison, we inferred an effective per-mutation failure rate from the observed data. With 174 mutations at 97% identity (for a 5.8kb genome), and assuming mutations contribute independently, the observed 5.3% viability implies:

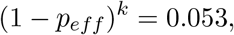

yielding *p*_*eff*_ = 1.67% ≈ (*CI* : 1.45 − 2.04%). This represents a summary statistic rather than a literal per-site lethality rate. The independence assumption is almost certainly wrong: epistasis is pervasive within the ΦX174 genome, which contains multiple overlapping reading frames, whilst the effects of mutation are not multiplicative in practice as we assume here. Therefore, the specific values we report should be treated as approximations, as epistatic interactions mean some mutation combinations are more or less deleterious than the product of individual effects. Calculating more accurate values would require significant dedicated experimental work beyond the scope of this manuscript. However, this effective rate provides a useful basis for comparison with literature values derived under similar assumptions, and the observed uplift is directionally real and plausibly accurate to an approximate order of magnitude.

#### Comparison to random mutagenesis

Deep mutational scanning of ΦX174 found that approximately 20% of random single mutations are lethal (Domingo-Calap et al., 2009). Importantly, these mutations were introduced randomly across the genome, which within a genome comprising 96% coding regions fell almost exclusively within such regions. Under this rate, the expected viability at 97% identity would be (0.8)^174^ ≈ 10^*−*17^ which is effectively zero. The model’s 5.3% viability therefore represents an uplift of many orders of magnitude over random mutagenesis (Figure 3B–C). In theory, the model might preferentially introduce synonymous mutations, which are largely neutral. Model uplift was not formally decomposed by mutation class; the difference in synonymous mutation rate between the model and random baselines was not sufficient to explain the observed degree of uplift.

Importantly, these rates measure different things: Domingo-Calap et al. (2009) measured lethality of random mutations; our *p*_*eff*_ reflects mutations the model chose to introduce. The difference could arise from site selection (avoiding critical regions), reduced individual deleteriousness (conservative substitutions), or favourable epistasis. It is possible that all three factors contribute.

#### Comparison to MSA-aware mutagenesis

A more generous baseline restricts mutations to sites that vary in natural Φ X174 sequences, where evolution has already demonstrated tolerance of substitution. Even under this assumption, achieving 5.3% viability would require implausibly low lethality rates at variable sites (*<* 0.5%). Across realistic rates (1–15%), the model still shows a substantial uplift of several orders of magnitude (Figure 3D). Design efficiency therefore cannot be explained solely by mutating at evolutionarily permissive positions.

#### Implications for genome length

When considering novel genome designs, efficiency might be expected to be increasingly consequential as genome length increases. Because, assuming constant lethality rates, viability under random mutation decays exponentially with length *P* ∝ (1 − *p*)^*L*^, even a modest per-site improvement would translate to large absolute gains for longer sequences (Figure 3E).

However, this extrapolation should be interpreted cautiously. ΦX174 is a 5.8kb ssDNA phage with extensively overlapping reading frames. This is an unusually constrained architecture in which almost the entire genome would be expected to be under selection. Larger genomes may have different constraint densities: some accessory or conditionally essential regions would be expected to be more tolerant to mutation, while others (regulatory elements, protein-protein interfaces) may be equally or more constrained. The relationship between genome length and mutational tolerance is unlikely to be uniform across viral families, and the exponential viability decay modelled here *P* ∝ (1−*p*)^*L*^ represents a worst case that assumes uniform genomic density of constraint.

We therefore present the genome length analysis as illustrative of how efficiency gains compound, rather than as a quantitative prediction for larger or more complex viruses. This framework would be most readily applicable to compact ssDNA and ssRNA viruses with little accessory or non-coding content, including bacteriophages such as ΦX174 and MS2, or eukaryotic viruses such as SARS-CoV-2.

Extending this framework to human-tropic pathogens would require empirical validation data from those systems, which, for obvious reasons, does not exist.

#### Comparison to serial passage

Serial passage is a powerful non-computational approach for improving viability, host range, or fitness through iterative selection. However, it explores sequence space locally and incrementally, typically requiring many generations to traverse larger mutational distances. Comparison with serial passage therefore provides a stringent evolutionary baseline: it asks whether generative models merely recapitulate what adaptive evolution can already achieve, or whether they confer a distinct efficiency advantage. Using typical serial passage simulation parameters (see Methods), we find that reaching comparable levels of evolutionary novelty requires impractically large numbers of passages (Fig. 3F), even though some trajectories may achieve higher fitness. A sensitivity analysis across 972 parameter combinations (Fig. 3G) shows that many scenarios fail to reach 97% sequence identity even after 2000 passages, corresponding to multiple years of daily passaging. These results indicate that generative genome design can achieve comparable exploratory outcomes with substantially greater time and cost efficiency than serial passage. Serial passage thus represents a baseline firmly in the Imitation quadrant - relatively inefficient and evolution-constrained, confirming the Evo 2 model-scaffold is not in the same quadrant.

#### Summary and framework placement

The Evo 2 model-scaffold achieves design efficiency exceeding many orders of magnitude over plausible baselines, not by exploring novel sequence space, but by effectively navigating the narrow viable region within evolutionary constraints. This confirms placement in the Optimisation quadrant: high efficiency, low novelty. The model substantially outperforms both random baselines and simple heuristics, but does so by staying close to what evolution has already explored.

## 3 Discussion

Our analysis provides a structured framework for evaluating generative genome design models by separating evolutionary novelty from design efficiency. Applying this framework to Evo 2-designed bacteriophages reveals a nuanced capability profile: the model functions as a highly efficient evolutionary optimiser rather than an unconstrained creative engine, placing it within the Optimisation quadrant.

### Evolutionary novelty: interpolation, not extrapolation

The model’s outputs remain phylogenetically close to natural sequences. Patristic distance, sequence identity, and genomic architecture all indicate that designed genomes cluster within existing evolutionary variation rather than representing novel clades. This suggests Evo 2 is largely interpolative: it recombines and reweights patterns present in training data,performing what might be called ‘semantic recombination’ of gene blocks in valid arrangements, rather than extrapolating toward fundamentally new biological designs.

### Design efficiency: substantial but bounded

Despite this evolutionary constraint, Evo 2 exhibits substantial design efficiency. The observed 5.3% viability at 97% identity corresponds to an effective per-mutation failure rate of approximately 1.7% - substantially lower than the 20% rate observed for random mutations in this phage. Compared to both naïve and MSA-aware baselines, the model enriches for viable sequences by many orders of magnitude. Importantly, this efficiency extends beyond what simple biological heuristics can explain: Evo 2 likelihood adds significant predictive value over covariates like GC content, gene synteny, and evolutionary similarity.

### Biosecurity implications

This evaluation was conducted on bacteriophages - a system chosen because it permits empirical validation with minimal biosecurity risk. The findings presented here are most readily applicable to other viruses that are similarly constrained, such as compact ssDNA and ssRNA viruses with genomes densely packed with coding sequences. They should not be straightforwardly extrapolated to large dsDNA phages or large eukaryotic DNA viruses. Indeed, the relationship between sequence novelty and function will differ across viral families, and ΦX174, with its extensively overlapping reading frames and almost non-existent non-coding regions, might well create a landscape of constraint that is not representative of larger viral genomes with substantial accessory content of conditional importance. What this analysis provides is a framework and proof-of-concept. We demonstrate that evolutionary novelty and design efficiency can be empirically assessed when functional validation data exists, and show how to locate a model-scaffold within this two-axis space. For Evo 2 on bacteriophages, it appears to be located within the Optimisation quadrant.

What does Optimisation quadrant-placement mean for biosecurity? High design efficiency within evolutionary constraints accelerates existing engineering workflows - reducing iteration time, lowering expertise barriers, and increasing throughput. This is not a trivial concern: making known, or subtly novel, biology easier to engineer has real implications for access. However, with the caveat that functional and phenotypic novelty would not be expected to scale linearly with phylogenetic distance, it does not represent the qualitatively novel capability that would come from reliable generation of functional sequences in unexplored regions of sequence space.

The key surveillance question for future models is whether they begin showing viability at greater phylogenetic distance - movement along the x-axis toward the Invention quadrant. Our framework provides a template for detecting such transitions.

## Limitations

Several limitations warrant consideration.

First, we analysed a dataset that had already been filtered for biological plausibility, including minimum spike protein homology. Our efficiency estimates therefore reflect a model-scaffold (model plus filtering) rather than raw model output. This is appropriate for assessing real-world capability; users will apply similar filters, but this means raw model performance may be lower than reported.

Second, our quantitative comparisons rely on an independence assumption that is almost certainly inaccurate. The lethality rate was constructed from Domingo-Calap et al, an analysis which measured the lethality rate of single mutations, whereas each genome assessed here had many novel mutations; within a genome as constrained as ΦX174, any assumption of mutation independence will inevitably be inaccurate. Epistatic interactions mean the effective per-mutation failure rate is a summary statistic rather than a literal per-site probability. More accurate assessments of combinatorial lethality rates would require experiments that are beyond the scope of this work. Furthermore, this baseline did not distinguish between synonymous and nonsynonymous mutations, and it is plausible that this rate differed compared to variation generated by the model. This effect could not plausibly account for the vast uplift demonstrated here. Nonetheless, we hope our simplification is transparent and illustrative.

Third, we characterised one model on one system at one identity threshold. ΦX174 is a small, highly constrained phage with overlapping reading frames, not representative of all viral architectures. Extending this framework to other systems, particularly larger or more complex genomes, would require additional empirical validation. Likewise, this analysis was conducted using a particular configuration of the model, with temperature, and post-hoc filtering stringency pre-defined. A more robust assessment of the place of this model in our framework would incorporate alternate sampling approaches.

Fourth, while we assessed the x-axis qualitatively through phylogenetic analysis, we did not establish a quantitative threshold for what constitutes “high” versus “low” novelty. Operationalising the framework more precisely remains an area for future development.

Finally, our framework requires functional validation data to locate models empirically. Such data is rarely available for generative genome design, and would be inadvisable to generate for pathogens of pandemic concern. This constrains where the framework can be directly applied, though the conceptual decomposition remains useful for structuring risk discussions even absent empirical validation.

## Conclusion

Our two-axis framework, evolutionary novelty and design efficiency, provides a transparent and generalisable approach for assessing generative genome models. For compact viral genomes, current systems, including Evo 2 behave more like powerful evolutionary priors than creative design engines: highly efficient within the bounds of known biology, but not yet capable of reliable invention beyond it. We do not offer evidence as to whether this characterisation extends to other, structurally distinct forms of genome, or subsections thereof. As models continue to advance, this framework offers a principled basis for evidence-based capability assessment and biosecurity evaluation.

## 4 Methods

### 4.1 Dataset Curation and Reference Genomes

We obtained the synthetic bacteriophage genomes generated by King et al. (2025). This dataset comprised 302 synthetic sequences generated by variants of the Evo genome language models that had been fine-tuned on naturally occurring bacteriophage sequences.

To derive ground truth results for sequence viability, we classified these sequences based on the *in vitro* validation results reported by King et al. (2025). We harmonised the list of generated sequences with list of validated sequences, confidently reconciling two validated sequences that lacked exact identifier matches by aligning them based on sequence length and identifier similarity. Furthermore, two sequences labelled ‘R1’ and ‘R2’ were excluded from our analysis, as they were not direct generative outputs of the model. Sequences were designated as ‘viable’ if they successfully produced plaques in the host bacterium, and ‘non-viable’ otherwise.

### 4.2 Generative Model Scoring

To evaluate the predictive capacity of genome language models for viral sequence viability, we scored all 302 synthetic sequences using three distinct architectures. We utilised: the 131k context length version of Evo1; the 7B parameter Evo 2 base model; the specific fine-tuned Evo 2 checkpoint developed by King et al. (2025) for bacteriophage design; the 250M parameter version of Nucleotide Transformer from Dalla-Torre et al. (2025), selected to represent an alternative class of genome language models, trained with a masked language modelling objective.

Sequence likelihoods were calculated as the joint probability of the nucleotide sequence under the Evo 2 models. For the Nucleotide Transformer, we utilised the pseudo-likelihood scoring method (Salazar et al., 2020), summing the conditional log-probabilities of each token when masked individually. These scores were subsequently used to generate Receiver Operating Characteristic (ROC) curves and calculate the Area Under the Curve (AUC) to assess the ability of each model to discriminate between viable and non-viable designs.

### 4.3 Phylogenetic Analysis and Evolutionary Distance

For comparative phylogenetic analysis, we assembled the same set of 11 naturally occurring reference genomes utilised by King et al. King et al. (2025) in their construction of phylogeny. Expanding this suite of genomes could refine future estimates of phylogenetic distance.

To quantify the evolutionary novelty of the synthetic designs, we examined their phylogenetic context relative to the natural reference genomes. We performed a multiple sequence alignment (MSA) of the 302 synthetic sequences and the 11 reference genomes using MAFFT (Katoh & Standley, 2013). Following alignment, we reconstructed a maximum-likelihood phylogenetic tree using IQ-TREE (Nguyen et al., 2015).

We calculated the “patristic distance” for each synthetic sequence, defined as the sum of branch lengths connecting the sequence node to the nearest natural reference node in the tree (Fourment & Gibbs, 2006). This metric served as a proxy for evolutionary divergence, allowing us to correlate model likelihoods with the degree to which sequences adhered to existing evolutionary patterns.

### 4.4 Biophysical and Structural Covariates

To assess the biological plausibility of the generated sequences, we computed a comprehensive suite of low-complexity covariates reflecting genomic stability, composition, and structural integrity.

- Global Sequence Identity (BLAST): To quantify true evolutionary novelty, we calculated the nucleotide identity of each synthetic sequence against the entire NCBI nucleotide database (nt) using BLASTn (Camacho et al., 2009). The maximum percent identity to the nearest match found globally serves as a proxy for the distance of that sequence relative to naturally occurring sequences.
- GC Content and Homopolymers: We calculated the global GC content for each synthetic genome and derived the absolute percentage difference relative to the wild-type ΦX174 reference genome. Additionally, we scanned sequences for homopolymeric runs, flagging instances of a single nucleotide repeating for multiple bases (such repeats can impede synthesis and replication King et al. (2025)).
- Genome Length Deviation: We calculated the absolute difference in nucleotide length between each synthetic candidate and the wild-type ΦX174 genome (5,386 bp) to quantify expansion or contraction of the genomic architecture.
- Gibson Assembly Thermodynamics (*T*_*m*_): To approximate the thermodynamic stability of the DNA fragments utilised for assembly, a metric reflecting the melting temperatures of the specific 5’ and 3’ junction overlaps was obtained from the King et al supplement. These *T*_*m*_ values were derived based on the nearest-neighbour thermodynamic model (SantaLucia, 1998) assuming standard PCR salt concentrations (Na^+^ = 50 mM, Mg^2+^ = 1.5 mM).
- Coding Density and ORF Counts: We defined coding density as the proportion of the total genome length occupied by predicted Open Reading Frames (ORFs). ORFs were predicted using Prodigal (in *meta* mode) (Hyatt et al., 2010). The raw count of predicted ORFs and the coding density were leveraged as distinct covariates.
- Protein Similarity (MMseqs2): To quantify proteomic divergence, predicted amino acid sequences were aligned against the proteome of the nearest phylogenetic neighbour using MMseqs2 (sensitivity setting 4.0) (Steinegger & Söding, 2017). The average percent identity of the top hits was calculated for all identified coding sequences.
- Syntenic Gene Count: The number of genes retaining conserved synteny per sequence was calculated. A gene was classified as syntenic if it appeared in the same relative transcriptional order as its homologue in the ΦX174 reference genome, as determined by pairwise proteomic alignment King et al. (2025).
- Architecture Similarity Score: This score was leveraged from the King et al. (2025) supplement. Briefly, the preservation of global genomic structure was estimated using a vectorised boundary scoring method. Genomic architectures were represented by one-hot encoding the positions of all start and stop codons. These vectors were subjected to Gaussian blurring (*σ* = 5) to allow for positional flexibility, and the similarity was computed as the dot product between the synthetic and reference architecture vectors, normalised to the self-similarity score of the wild-type ΦX174 genome King et al. (2025).
- Mapping Proportion: We aligned each synthetic genome to the ΦX174 reference using Minimap2 (Li, 2018). The “Mapping Proportion” is reported as the percentage of the synthetic genome length that successfully aligns to the reference, serving as a proxy for global structural coherence.

### 4.5 Quantification of Generative Uplift

To separate the model’s contribution from simple evolutionary conservation, we quantified ‘Design Efficiency’ as the increase in the probability of generating a viable sequence relative to a random baseline.

First, we inferred an effective per-mutation failure rate for the model. We observed a viability rate of 5.3% for genomes generated at approximately 97% nucleotide identity. Assuming an independent-hits model, we calculated the effective per-mutation failure probability (*p*_eff_) using the equation:

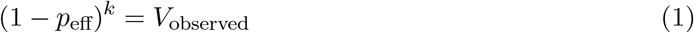

where *k* is the number of mutations (174 for a 5.8kb genome) and *V*_observed_ is the empirical viability (0.053).

We then calculated the expected viability of baseline generators (*P*_baseline_) under two scenarios:

1. Naïve Baseline: Assumes mutations are distributed uniformly with a per-mutation lethality rate of 20% (*p*_lethal_ = 0.2), based on deep mutational scanning data (Domingo-Calap et al., 2009).
2. MSA-Aware Baseline: Restricts mutations solely to sites identified as variable within the multiple sequence alignment, testing across a range of assumed lethal rates (1–15%).

We define Generative Uplift as the ratio of the model’s viability probability to the baseline viability probability at a given number of mutations *k*:

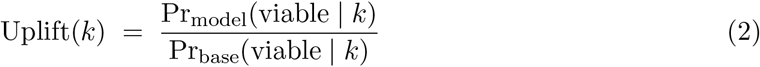

This general formulation allows comparison against any baseline generator. For the naïve random mutagenesis baseline, where each mutation has independent lethality probability *p*_lethal_, this becomes:

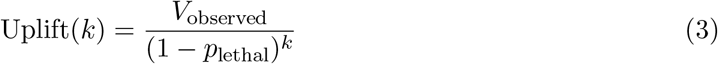

For the MSA-aware baseline, we substitute the assumed lethality rate at variable sites (*p*_var_) for

*p*_lethal_, testing across a range of plausible values (0.1–15%).

### 4.6 Statistical Analysis

All statistical analyses were performed using R (version 4.2.2). Data wrangling and visualisation were conducted using tidyverse alongside ggpubr and gridExtra.

Receiver Operating Characteristic (ROC) analysis was performed using the pROC package to calculate AUC metrics. We first screened potential predictors of viability using univariate logistic regression. Variables that demonstrated statistical significance (*p <* 0.05) in this univariate screening were subsequently included in a multivariable logistic regression model. Model outputs were processed using the broom package.

To quantify the added value of the Evo 2 model, we compared a baseline model (comprising the significant low-complexity covariates) against a full model (baseline + Evo 2 likelihoods). We assessed the improvement in model fit using the Likelihood Ratio Test and calculated McFadden’s pseudo-*R*^2^ to quantify the proportion of variance explained by each model configuration.

### 4.7 Serial-passage evolutionary simulations

We simulated viral evolution under repeated serial passage using a minimal stochastic population-genetic model capturing mutation, selection, and inter-passage bottlenecks. The model is designed to quantify how rapidly genome-wide sequence identity can diverge from an ancestral sequence under experimentally realistic conditions.

#### Model overview

Each viral genome is represented by the cumulative number of substitutions in three fitness classes: deleterious (*k*_*d*_), neutral (*k*_*n*_), and beneficial (*k*_*b*_). The total number of substitutions is *k* = *k*_*d*_ + *k*_*n*_ + *k*_*b*_. Fitness is assumed multiplicative:

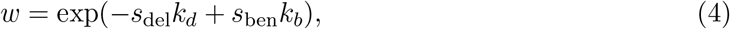

where *s*_del_ and *s*_ben_ are per-mutation selection coefficients.

Within each passage, the population evolves for *g* effective viral generations. Each generation consists of:

##### 1. Selection and drift

A Wright–Fisher (Wright, 1931) sampling step with population size *N*, where offspring are sampled multinomially with probabilities proportional to parental abundance weighted by fitness.

##### 2. Mutation

Each offspring genome acquires new point mutations with probability *λ* = *Lµ*, where *L* is genome length and *µ* is the per-site mutation rate per generation. For computational efficiency (valid because *λ* ≪ 1 for all parameter values used), at most one mutation per genome per generation is assumed. Mutations are assigned to deleterious, neutral, or beneficial classes with probabilities *p*_del_, *p*_neu_, *p*_ben_.

After *g* generations, a **bottleneck** is imposed by sampling *B* genomes multinomially from the population to seed the next passage. This captures experimental transfer, dilution, or plaque-picking effects. The process is repeated for *P* passages.

#### Sequence identity calculation

The simulator tracks cumulative substitutions per site:

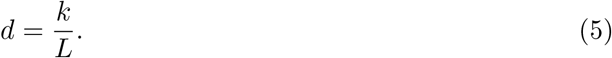

Observed sequence identity is derived using the Jukes–Cantor (JC69) substitution model to account for multiple hits and back mutations (Jukes et al., 1969):

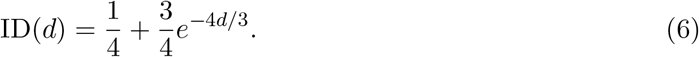

We report both the population-mean identity and a consensus-like identity computed from the most abundant genotype (modal lineage). All reported identities refer to the JC-corrected expected sequence identity.

#### Parameterisation

Unless otherwise stated, simulations were parameterised to approximate a small single-stranded DNA bacteriophage (ΦX174-like) genome:

- Genome length: *L* = 5,386 nt
- Total passages: *P* = 2,000
- Within-passage effective population size: *N* = 10^7^

We explored a grid of biologically plausible values for the dominant evolutionary control parameters.

#### Bottleneck size

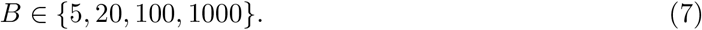

#### Effective generations per passage

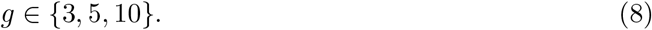

#### Per-site mutation rate

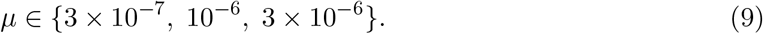

#### Fraction of deleterious mutations

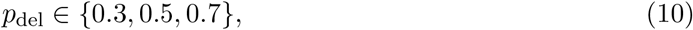

with *p*_ben_ = 10^*−*3^ fixed and

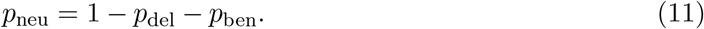

These values represent the effective fraction of mutations contributing to long-term substitution dynamics, incorporating genome-wide constraint, epistasis, and repeated bottlenecks.

#### Selection coefficients

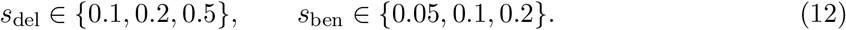

This grid yielded 972 distinct parameter combinations.

### Replicates and stochasticity

Each parameter combination was simulated with three independent random seeds. For each condition, we summarised median final sequence identity, quantiles where appropriate, and whether identity crossed a predefined threshold (97%).

Because our primary conclusions concern large-scale separation between parameter regimes rather than fine-grained stochastic variability, a small number of seeds was sufficient to establish robustness; representative subsets were additionally validated with higher replicate counts.

### Output metrics

For each simulation we recorded:

- Final population-mean sequence identity.
- Final consensus-like sequence identity (modal genotype).
- Trajectories of identity across passages.
- Time to reach predefined identity thresholds when applicable.

Unless otherwise stated, figures report final consensus-like sequence identity after 2,000 passages.

## Acknowledgments

JRMB and AM are funded by Coefficient Giving. OMC is funded by an MRC grant (MR/Y010078/1) and Todd-Bird research fellowship at the New College, Oxford. The authors thank Theo Sanderson, Damon Binder, Steph Guerra, Chris Barnes, Luca Righetti, Yunha Hwang, and Jeremy Ratcliff for feedback and discussions.

## Data and Code Availability

Analysis code and derived data (model likelihood scores, phylogenetic distances) are available on *GitHub* and archived at Zenodo (DOI: XXX). Primary sequence data and experimental validation results were obtained from King et al. (2025).

## References

Almeida, B. P. d., Dalla-Torre, H., Richard, G., Blum, C., Hexemer, L., Gélard, M., Mendoza-Revilla, J., Tang, Z., Marin, F. I., Emms, D. M., Pandey, P., Laurent, S., Lopez, M., Laterre, A., Lang, M., Şahin, U., Beguir, K., and Pierrot, T. Annotating the genome at single-nucleotide resolution with DNA foundation models, March 2025. URL https://www.biorxiv.org/content/10.1101/2024.03.14.584712v4. Pages: 2024.03.14.584712 Section: New Results.

Benegas, G., Batra, S. S., and Song, Y. S. DNA language models are powerful predictors of genome-wide variant effects. Proceedings of the National Academy of Sciences, 120(44): e2311219120, October 2023. ISSN 0027-8424, 1091-6490. doi: 10.1073/pnas.2311219120. URL https://pnas.org/doi/10.1073/pnas.2311219120.

Benegas, G., Albors, C., Aw, A. J., Ye, C., and Song, Y. S. A DNA language model based on multispecies alignment predicts the effects of genome-wide variants. Nature Biotechnology, 43 (12):1960–1965, December 2025. ISSN 1546-1696. doi: 10.1038/s41587-024-02511-w. URL https://www.nature.com/articles/s41587-024-02511-w. Publisher: Nature Publishing Group.

Black, J. R. M., Hanke, M. S., Maiwald, A., Hernandez-Boussard, T., Crook, O. M., and Pannu, J. Open-weight genome language model safeguards: Assessing robustness via adversarial fine-tuning, November 2025. URL http://arxiv.org/abs/2511.19299. arXiv:2511.19299 [cs].

Bloomfield, D., Pannu, J., Zhu, A. W., Ng, M. Y., Lewis, A., Bendavid, E., Asch, S. M., Hernandez-Boussard, T., Cicero, A., and Inglesby, T. AI and biosecurity: The need for governance. Science, 385(6711):831–833, August 2024. doi: 10.1126/science.adq1977. URL https://www.science.org/doi/10.1126/science.adq1977. Publisher: American Association for the Advancement of Science.

Brixi, G., Durrant, M. G., Ku, J., Poli, M., Brockman, G., Chang, D., Gonzalez, G. A., King, S. H., Li, D. B., Merchant, A. T., Naghipourfar, M., Nguyen, E., Ricci-Tam, C., Romero, D. W., Sun, G., Taghibakshi, A., Vorontsov, A., Yang, B., Deng, M., Gorton, L., Nguyen, N., Wang, N. K., Adams, E., Baccus, S. A., Dillmann, S., Ermon, S., Guo, D., Ilango, R., Janik, K., Lu, A. X., Mehta, R., Mofrad, M. R. K., Ng, M. Y., Pannu, J., Ré, C., Schmok, J. C., John, J. S., Sullivan, J., Zhu, K., Zynda, G., Balsam, D., Collison, P., Costa, A. B., Hernandez-Boussard, T., Ho, E., Liu, M.-Y., McGrath, T., Powell, K., Burke, D. P., Goodarzi, H., Hsu, P. D., and Hie, B. L. Genome modeling and design across all domains of life with Evo 2, February 2025. URL https://www.biorxiv.org/content/10.1101/2025.02.18.638918v1. Pages: 2025.02.18.638918 Section: New Results.

Camacho, C., Coulouris, G., Avagyan, V., Ma, N., Papadopoulos, J., Bealer, K., and Madden, T. L. BLAST+: architecture and applications. BMC Bioinformatics, 10(1):421, December 2009. ISSN 1471-2105. doi: 10.1186/1471-2105-10-421. URL https://doi.org/10.1186/1471-2105-10-421.

Dalla-Torre, H., Gonzalez, L., Mendoza-Revilla, J., Lopez Carranza, N., Grzywaczewski, A. H., Oteri, F., Dallago, C., Trop, E., de Almeida, B. P., Sirelkhatim, H., Richard, G., Skwark, M., Beguir, K., Lopez, M., and Pierrot, T. Nucleotide Transformer: building and evaluating robust foundation models for human genomics. Nature Methods, 22(2):287–297, February 2025. ISSN 1548-7105. doi: 10.1038/s41592-024-02523-z. URL https://www.nature.com/articles/s41592-024-02523-z. Publisher: Nature Publishing Group.

Domingo-Calap, P., Cuevas, J. M., and Sanjuán, R. The Fitness Effects of Random Mutations in Single-Stranded DNA and RNA Bacteriophages. PLOS Genetics, 5(11):e1000742, November 2009. ISSN 1553-7404. doi: 10.1371/journal.pgen.1000742. URL https://journals.plos.org/plosgenetics/article?id=10.1371/journal.pgen.1000742. Publisher: Public Library of Science.

Fishman, V., Kuratov, Y., Shmelev, A., Petrov, M., Penzar, D., Shepelin, D., Chekanov, N., Kardymon, O., and Burtsev, M. GENA-LM: a family of open-source foundational DNA language models for long sequences. Nucleic Acids Research, 53(2):gkae1310, January 2025. ISSN 1362-4962. doi: 10.1093/nar/gkae1310. URL https://doi.org/10.1093/nar/gkae1310.

Fourment, M. and Gibbs, M. J. PATRISTIC: a program for calculating patristic distances and graphically comparing the components of genetic change. BMC Evolutionary Biology, 6:1, January 2006. ISSN 1471-2148. doi: 10.1186/1471-2148-6-1. URL https://pmc.ncbi.nlm.nih.gov/articles/PMC1352388/.

Hatfull, G. F. and Hendrix, R. W. Bacteriophages and their Genomes. Current opinion in virology, 1(4):298–303, October 2011. ISSN 1879-6257. doi: 10.1016/j.coviro.2011.06.009. URL https://pmc.ncbi.nlm.nih.gov/articles/PMC3199584/.

Hyatt, D., Chen, G.-L., LoCascio, P. F., Land, M. L., Larimer, F. W., and Hauser, L. J. Prodigal: prokaryotic gene recognition and translation initiation site identification. BMC Bioinformatics, 11(1):119, March 2010. ISSN 1471-2105. doi: 10.1186/1471-2105-11-119. URL https://doi.org/10.1186/1471-2105-11-119.

Jha, N., Kravitz, J., West-Roberts, J., Camargo, A., Roux, S., Cornman, A., and Hwang, Y. Gaia: A Context-Aware Sequence Search and Discovery Tool for Microbial Proteins, November 2024. URL https://www.biorxiv.org/content/10.1101/2024.11.19.624387v1. Pages: 2024.11.19.624387 Section: New Results.

Ji, Y., Zhou, Z., Liu, H., and Davuluri, R. V. DNABERT: pre-trained Bidirectional Encoder Representations from Transformers model for DNA-language in genome. Bioinformatics, 37 (15):2112–2120, August 2021. ISSN 1367-4803. doi: 10.1093/bioinformatics/btab083. URL https://doi.org/10.1093/bioinformatics/btab083.

Jukes, T. H., Cantor, C. R., et al. Evolution of protein molecules. Mammalian protein metabolism, 3(21):132, 1969.

Katoh, K. and Standley, D. M. MAFFT Multiple Sequence Alignment Software Version 7: Improvements in Performance and Usability. Molecular Biology and Evolution, 30(4):772–780, April 2013. ISSN 0737-4038. doi: 10.1093/molbev/mst010. URL https://doi.org/10.1093/molbev/mst010.

Kilcher, S. and Loessner, M. J. Engineering Bacteriophages as Versatile Biologics. Trends in Microbiology, 27(4):355–367, April 2019. ISSN 1878-4380. doi: 10.1016/j.tim.2018.09.006.

King, S. H., Driscoll, C. L., Li, D. B., Guo, D., Merchant, A. T., Brixi, G., Wilkinson, M. E., and Hie, B. L. Generative design of novel bacteriophages with genome language models, September 2025. URL https://www.biorxiv.org/content/10.1101/2025.09.12.675911v1. xISSN: 2692-8205 Pages: 2025.09.12.675911 Section: New Results.

Li, H. Minimap2: pairwise alignment for nucleotide sequences. Bioinformatics, 34(18):3094– 3100, September 2018. ISSN 1367-4803. doi: 10.1093/bioinformatics/bty191. URL https://doi.org/10.1093/bioinformatics/bty191.

Mendoza-Revilla, J., Trop, E., Gonzalez, L., Roller, M., Dalla-Torre, H., De Almeida, B. P., Richard, G., Caton, J., Lopez Carranza, N., Skwark, M., Laterre, A., Beguir, K., Pierrot, T., and Lopez, M. A foundational large language model for edible plant genomes. Communications Biology, 7(1):835, July 2024. ISSN 2399-3642. doi: 10.1038/s42003-024-06465-2. URL https://www.nature.com/articles/s42003-024-06465-2.

Nguyen, E., Poli, M., Durrant, M. G., Kang, B., Katrekar, D., Li, D. B., Bartie, L. J., Thomas, A. W., King, S. H., Brixi, G., Sullivan, J., Ng, M. Y., Lewis, A., Lou, A., Ermon, S., Baccus, S. A., Hernandez-Boussard, T., Ré, C., Hsu, P. D., and Hie, B. L. Sequence modeling and design from molecular to genome scale with Evo. Science, 386(6723):eado9336, November 2024. doi: 10.1126/science.ado9336. URL https://www.science.org/doi/10.1126/science.ado9336. Publisher: American Association for the Advancement of Science.

Nguyen, L.-T., Schmidt, H. A., von Haeseler, A., and Minh, B. Q. IQ-TREE: A Fast and Effective Stochastic Algorithm for Estimating Maximum-Likelihood Phylogenies. Molecular Biology and Evolution, 32(1):268–274, January 2015. ISSN 0737-4038. doi: 10.1093/molbev/msu300. URL https://doi.org/10.1093/molbev/msu300.

Pannu, J., Bloomfield, D., MacKnight, R., Hanke, M. S., Zhu, A., Gomes, G., Cicero, A., and Inglesby, T. V. Dual-use capabilities of concern of biological AI models. PLOS Computational Biology, 21(5):e1012975, August 2025. ISSN 1553-7358. doi: 10.1371/journal.pcbi.1012975. URL https://journals.plos.org/ploscompbiol/article?id=10.1371/journal.pcbi.1012975. Publisher: Public Library of Science.

Ranveer, S. A., Dasriya, V., Ahmad, M. F., Dhillon, H. S., Samtiya, M., Shama, E., Anand, T., Dhewa, T., Chaudhary, V., Chaudhary, P., Behare, P., Ram, C., Puniya, D. V., Khedkar, G. D., Raposo, A., Han, H., and Puniya, A. K. Positive and negative aspects of bacteriophages and their immense role in the food chain. npj Science of Food, 8(1):1, January 2024. ISSN 2396-8370. doi: 10.1038/s41538-023-00245-8. URL https://www.nature.com/articles/s41538-023-00245-8.

Salazar, J., Liang, D., Nguyen, T. Q., and Kirchhoff, K. Masked Language Model Scoring. In Jurafsky, D., Chai, J., Schluter, N., and Tetreault, J. (eds.), Proceedings of the 58th Annual Meeting of the Association for Computational Linguistics, pp. 2699–2712, Online, July 2020. Association for Computational Linguistics. doi: 10.18653/v1/2020.acl-main.240. URL https://aclanthology.org/2020.acl-main.240/.

SantaLucia, J. A unified view of polymer, dumbbell, and oligonucleotide DNA nearest-neighborthermodynamics. Proceedings of the National Academy of Sciences, 95(4):1460–1465, February 1998. doi: 10.1073/pnas.95.4.1460. URL https://www.pnas.org/doi/10.1073/pnas.95.4.1460. Publisher: Proceedings of the National Academy of Sciences.

Schiff, Y., Kao, C.-H., Gokaslan, A., Dao, T., Gu, A., and Kuleshov, V. Caduceus: Bi-Directional Equivariant Long-Range DNA Sequence Modeling, June 2024. URL http://arxiv.org/abs/2403.03234. arXiv:2403.03234 [q-bio].

Steinegger, M. and Söding, J. MMseqs2 enables sensitive protein sequence searching for the analysis of massive data sets. Nature Biotechnology, 35(11):1026–1028, November 2017. ISSN 1546-1696. doi: 10.1038/nbt.3988.

Wei, B., Che, Z., Li, N., Sehwag, U. M., Götting, J., Nedungadi, S., Michael, J., Yue, S., Hendrycks, D., Henderson, P., Wang, Z., Donoughe, S., and Mazeika, M. Best Practices for Biorisk Evaluations on Open-Weight Bio-Foundation Models, November 2025. URL http://arxiv.org/abs/2510.27629. xarXiv:2510.27629 [cs].

Wright, S. Evolution in mendelian populations. Genetics, 16(2):97, 1931.

Zhai, J., Gokaslan, A., Schiff, Y., Berthel, A., Liu, Z.-Y., Lai, W.-Y., Miller, Z. R., Scheben, A., Stitzer, M. C., Romay, M. C., Buckler, E. S., and Kuleshov, V. Cross-species modeling of plant genomes at single-nucleotide resolution using a pretrained DNA language model. Proceedings of the National Academy of Sciences, 122(24):e2421738122, June 2025. doi: 10.1073/pnas.2421738122. URL https://www.pnas.org/doi/10.1073/pnas.2421738122.

